# Chunking sequence information by mutually predicting recurrent neural networks

**DOI:** 10.1101/215392

**Authors:** Toshitake Asabuki, Naoki Hiratani, Tomoki Fukai

## Abstract

Interpretation and execution of complex sequences is crucial for various cognitive tasks such as language processing and motor control. The brain solves this problem arguably by dividing a sequence into discrete chunks of contiguous items. While chunking has been accounted for by predictive uncertainty, alternative mechanisms have also been suggested, and the mechanism underlying chunking is poorly understood. Here, we propose a class of unsupervised neural networks for learning and identifying repeated patterns in sequence input with various degrees of complexity. In this model, a pair of reservoir computing modules, each of which comprises a recurrent neural network and readout units, supervise each other to consistently predict others’ responses to frequently recurring segments. Interestingly, this system generates neural responses similar to those formed in the basal ganglia during habit formation. Our model extends reservoir computing to higher cognitive function and demonstrates its resemblance to sequence processing by cortico-basal ganglia loops.

## Introduction

When a sequence of stimuli is repeated, these stimuli may be segmented and then bound together into “chunks” that are stored and processed as single units. Chunking or “bracketing” (Graybiel, 1998) appears in various cognitive behaviors that require sequence processing (Miller, 1956; Ericcson et al., 1980; Sakai et al., 2004; Orban et al., 2007; Christiansen and Chater, 2016). For instance, in language acquisition continuous vocal sounds are segmented into recurring groups of contiguous sounds that are processed as words (Builatti et al., 2009; Estes et al., 2007; Gentner et al., 2006). A sequence of movements may be executed as one compound movement after repetitive practice, which is thought to be the formation of motor habits (Fujii and Graybiel, 2003; Graybiel 1998; Jin et al., 2014; Smith and Graybiel, 2013). In the context of sequence generation, chunking is thought to reduce the complexity of sequence processing and the associated cost (Miller 1956; Ramkumar et al., 2016; Verwey and Abrahamse, 2012). Thus, chunking constitutes a crucial step in representing the hierarchical structure of sequential knowledge (Dehaene et al., 2015).

However, chunking is still a challenge in neural computation. Chunking has been thought to occur through two processes. Long and complex sequences are first segmented into shorter and simple sequences. Then, frequently repeated segments may be concatenated into a single unit (Wymbs et al., 2012). Various mechanisms of chunking have been proposed based on Bayesian computation (Orban et al., 2007; Kiebel et al., 2009), statistical learning guided by prediction errors (Reynolds et al., 2007), a bifurcation structure in nonlinear dynamical systems (stable heteroclinic orbits) (Rabinovich et al., 2014; Fonollosa et al., 2015), and even a neuromorphic hardware has been proposed (Guoqi et al., 2016). However, whether a bifurcation theoretic mechanism enables flexible chunking of complex sequences remains elusive and a recent experiment, as explained below, has suggested a prediction-free mechanism (Schapiro et al., 2013). Therefore, the neural mechanisms of flexible chunk formation remain unclear.

A widely accepted hypothesis is that chunking relies on prediction errors or surprise. Evidence supporting this hypothesis is typically obtained from studies of mismatch negativity, a brain signal (such as electoencepharogram and functional magnetic resonance imaging) indicating the detection of deviance from a regular temporal pattern of sensory stimuli (Bekinschtein et al., 2009; Chait et al., 2012; Schröger et al., 2014; Wacongne et al., 2012; Uhrig et al., 2014; Näätänen, 2003; Garrido et al., 2009; Kremláček et al., 2016). It is likely that deviant stimuli are detectable if the brain has a prediction on the stimulus pattern. In other words, the brain should know the recurring sequence patterns before it can perceive deviance. Actually, the brain can detect the repetition of patterns in random sequences (Huettel et al., 2002; Romberg and Saffran, 2013), and it is likely that a common neural mechanism may underlie chunking and such a pattern detection. In fact, chunking favors an account based on the temporal community detection, in which the stimuli that frequently go together are grouped into a chunk (Schapiro et al., 2013). However, the underlying mechanism of those cognitive functions largely remains unclear.

In this study, we propose a novel mechanism of unsupervised chunk learning without relying on prediction errors. To this end, we utilize a recurrent network model for cortical computation (Maass et al, 2002; Jaeger and Haas, 2004). We extend the framework of reservoir computing (RC) to unsupervised learning. RC consists of a recurrent neural network, readout units, and feedforward and feedback projections between them, and undergoes supervised learning in its original form (Sussillo and Abbott, 2009). The key of our proposal is to use a pair of independent RC modules that supervise each other. Sequence leaning with RC has been extensively studied in motor control (Laje and Buonomano, 2013; Shenoy et al., 2011; Sussillo et al., 2015) and decision making (Mante et al., 2013; Carnevale et al., 2015), and theoretical extensions have also been proposed, for instance, to spiking neuron networks (Abbott et al., 2016) and/or reward-based learning (Hoerzer et al., 2014). In our model, teaching signals necessary for supervised learning are provided by the partner networks, and consequently the entire RC system learns in an unsupervised fashion to extract various irregularly recurring patterns in complex sequences.

Another biological implication of our model is that it self-organizes task-related activities similar to those formed in the basal ganglia during motor habit formation. Some striatal neurons respond selectively to the first (Start cells) or the last (Stop cells), or both, element of a motor sequence after a repetitive training of sequence execution (Jin et al., 2014; Smith and Graybiel, 2013). Because all the elements of the sequence are equally represented in the striatum before training, the Start/Stop cells are thought to encode the motor chunks acquired by the training. Our model consistently replicates task-related neurons similar to the Stop cells through learning an arbitrary complex sequence.

## RESULTS

### Reservoir computing modules with mutual supervision

To demonstrate the basic framework of our model, we first consider the case where input sequence alternates a single chunk (i.e., a-b-c-d) and random sequences of discrete items. To be specific, we use 26 letters of the English alphabet (e to z) to denote these items (Fig. 1a). In reality, each alphabet may correspond to a brief stimulus in any sensory modality such as a brief tone signal. The random sequence components are introduced to unambiguously define the initial and end points of a chunk, and their lengths vary in every repetition cycle within the length range of 5 to 8. Each alphabet causes a phasic activation of synaptic current from the corresponding input neuron with slow rise and decay constants (Fig. 1b).

**Figure 1.**
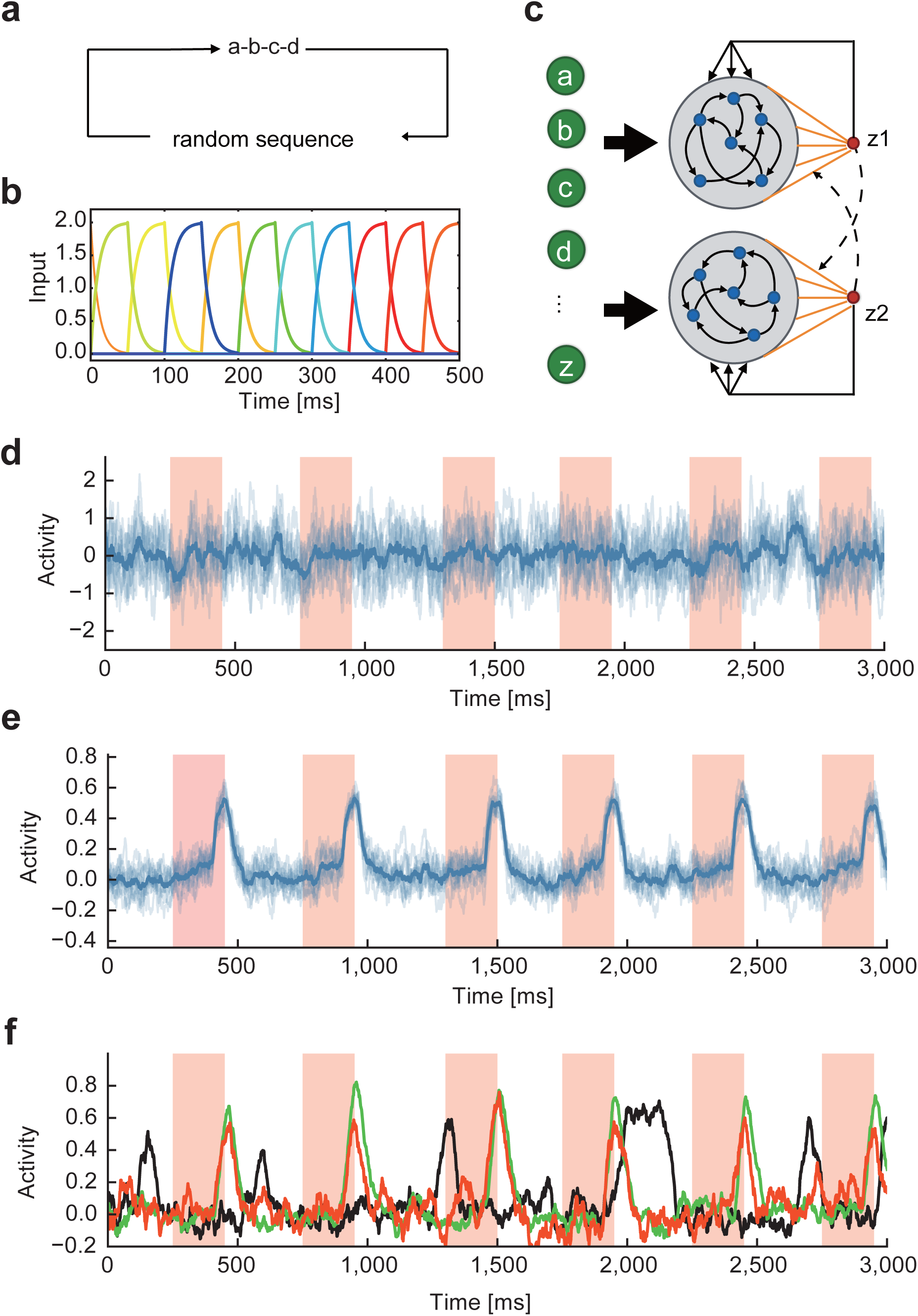
Learning of a single chunk repeated in random sequence. **(a)** Input sequence repeated a single chunk. In this example, chunk is composed of four alphabets (a, b, c, d). The components and lengths of random sequences varied during the repetition of chunks. **(b)** Example responses are shown for input neurons. **(c)** In the dual RC model, two non-identical reservoirs are activated by the same set of input neurons. Readout weights of each RC system undergo supervised learning with a teaching signal given by the output of the partner network. **(d) and (e)** Pre- and post-learning trial averaged activities of a readout unit are shown, respectively. Shaded intervals designate the presentation periods of the chunk. The other readout unit exhibited a similar activity pattern. **(f)** Readout activity was trained with many-to-one input projections. The fraction of input neurons projecting to a reservoir neuron was 10% (red), 40% (green) and 70% (black).

Our network model comprises two mutually non-interacting RC modules, each of which consists of a recurrent network (reservoir) of rate-based neurons and a readout unit, and receives an identical input sequence (Fig. 1c). Each reservoir neuron receives a selective input from one of the input neurons. However, this condition is not essential for chunking and can be relaxed, as shown later. Within each reservoir, all neurons are mutually connected and project to a readout unit, which projects back to all neurons belonging to the same reservoir. Note that the two reservoirs have different recurrent wiring patterns and hence are not identical. Activity of each readout unit *z(t)* is given as a weighed sum of the activities ***r***(*t*) of reservoir neurons projecting to the readout: *z(t)* = ***w***^T^***r***(*t*). Note that one readout unit per reservoir is sufficient for learning a single chunk. We will consider more complex cases later. The weight vector ***w*** is modifiable through the FORCE learning algorithm (Sussillo and Abbott, 2009), whereas recurrent and feedback connections are non-plastic because the model can solve the present task without modifying these connections. The initial states of the reservoirs are weakly chaotic as in the previous model (Sussillo and Abbott, 2009). See the Methods for the details of the model and the values of parameters.

A unique feature of the present model is that the output of each readout unit is used as a teacher signal to train the readout weights of the other reservoir module, implying that the two RC modules supervise each other. As a consequence, though the FORCE learning per se is a supervised learning rule, the entire network, which we may call “dual RC system”, is subject to unsupervised leaning because teaching signals originate from the system itself. The details of the teaching signals will be shown later.

### Chunk learning from a random sequence

The design of teaching signals is the key for successful chunk learning in the present model. The teaching signals should be symmetric with respect to the interchange of the two readout units, and should be determined such that the two systems stop learning when the two readout units output similar response patterns. In other words, the teaching signals eventually become identical between the two RC modules during the learning. The following teaching signals *f*_*i*_ enable chunk learning in the proposed dual RC system:

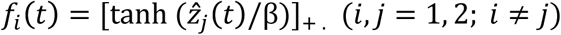

where 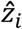 is the normalized output of the *i*-th readout unit (Methods), the threshold linear function [*x*]_+_ returns 0 if *x*≦0, and [*x*]_+_ = *x* if *x*>0, and the constant was set as *β* = 3. Defining error signals as *e*_*i*_(*t*) = *z*_*i*_(*t*) − *f*_*i*_(*t*), we train the pair of RC modules through the FORCE learning algorithm until the error signals become sufficiently small (typically, about 0.01) and the readout weights converge to equilibrium values (within small fluctuations). Note that the sigmoidal function allows the system to learn nontrivial solutions *Z*_*j*_(*t*) ≠ 0, and also maintains the outputs, hence the teaching signals, finite during learning. The threshold linear function makes the outputs positive. These nonlinear transformations greatly improved the performance of learning. Importantly, the teaching signals do not explicitly contain information about the structure and timing of chunks in input sequence.

This dual RC system converged to a state of stable operations when the two RC systems produced similar teaching signals (hence similar outputs) that were consistent with the temporal structure of input sequence (Supplementary Fig. 1). The readout units didn’t respond to the chunk before learning (Fig. 1d). After learning, the responses of the readout units were tested for the input sequences that had not been used for the training. The test sequences contained the same chunk “a-b-c-d”, but the random sequence part was different. The readout units exhibited steady phasic responses time-locked to the chunk (Figs. 1e). The readout activity piled up gradually in the beginning of the chunk, rapidly increased at its end, and then returned to a baseline level outside of it. The selective responses to the chunk was also successfully learned when each reservoir neuron was innervated by multiple input neurons. As shown in Fig. 1f, the system succeeded in learning when 10% and 40% of input neurons projected to a reservoir neuron, but failed when the fraction was 70%. Thus, the responses of individual reservoir neurons should be sufficiently independent of each other to robustly capture the recurrence of chunks.

### Learning of multiple chunks

We can extend the previous learning rule for the learning of multiple chunks without much difficulty. To show this, we now embedded three chunks into a random input sequence (Fig. 2a). The three chunks had the same occurrence probability of 1/3. To process this complex input sequence, we made two modifications in the previous model. First, each reservoir was now connected to three readout units (*z*_1_, *z*_2_, *z*_3_ for the 1st reservoir and *z*_4_, *z*_5_, *z*_6_ for the 2nd reservoir) each responsible for the learning of one of the three chunks (Fig. 2b). Second, we modified teaching signals as follows:

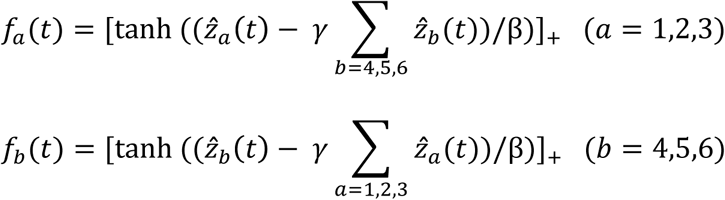

**Figure 2.**
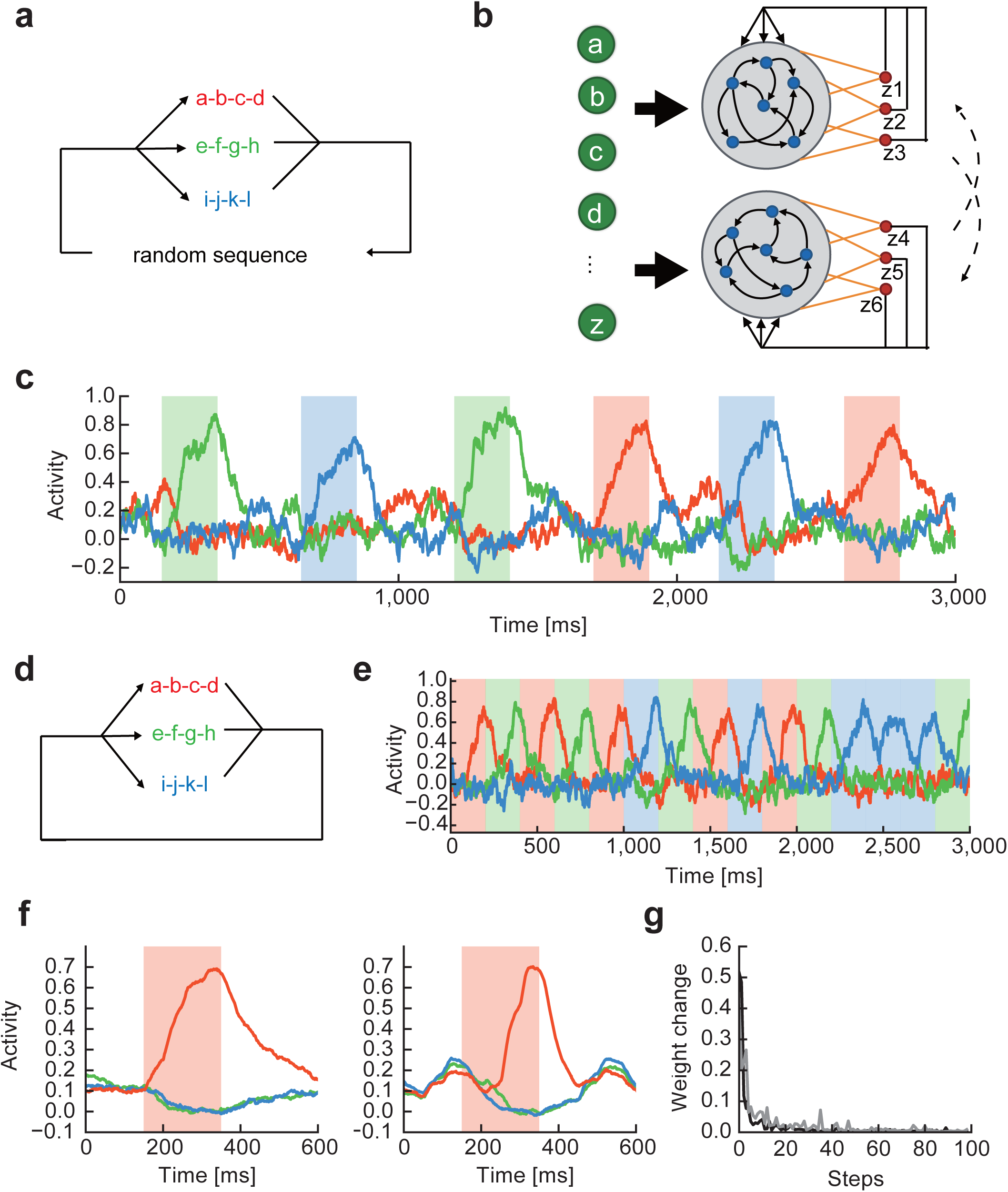
Readout activity after learning detects multiple chunks. **(a)** In input, three chunks a-b-c-d (red), e-f-g-h (green), and i-j-k-l (blue) separated by random sequences recurred at equal frequencies. **(b)** Each reservoir was connected to three readout units. **(c)** Selective readout responses to the individual chunks (colored intervals) were self-organized. The responses are colored according to their selectivity to the chunks. **(d)** The same chunks were repeated without the intervals of random sequences. Previous models of chunking typically processed such input sequences. **(e)** Selective readout responses were formed for the individual chunks. **(f)** Readout activities formed with (left) and without (right) random sequence intervals were averaged over the recurrence of chunk “a-b-c-d”. **(g)** Time evolution of average readout weights is shown at every 50 [s], which corresponds to “one step” in the abscissa, during learning for trials with (gray) and without (black) random sequence intervals.

where constant *γ* was set as 0.5 and the indices represent the exchanges of teaching signals between the RC modules. These teaching signals are symmetric with respect to the permutation of indices per reservoir and allow each output unit to adopt to a specific chunk. A further extension of the learning rule to an arbitrary number of chunks is straightforward.

As in the case with a single chunk, all readout units displayed a ramping activity selective to a specific chunk, signaling a successful chunk learning (Fig. 2c). The question then arises whether the RC system could learn the multiple chunks as they were temporally separated by random sequences. To show this is not the case, we trained the model by using input sequences in which three chunks appear randomly and consecutively with equal probabilities without any interval of random sequences (Fig. 2d). The same RC system as before could easily learn the multiple chunks (Fig. 2e). A notable difference was that outside of the chunks the readout activity decayed faster for the undisturbed sequences than for the temporally separated ones (Fig. 2f). In fact, learning proceeded a little faster for the former sequences (Fig. 2g), suggesting that the learning is more effective when chunks are not disrupted by random sequences.

### Selective recruitment of reservoir neurons for chunk learning

Next, we investigate how the activities of reservoir neurons encode chunks. Here, the network was trained on sequences containing three chunks and random sequences. In each reservoir, a subset of neurons selectively responded to each chunk after learning (Fig. 3a). We therefore classified reservoir neurons into three ensembles according to the selectivity of their responses to each chunk (Materials and Methods). Although some reservoir neurons responded to more than one chunk, we excluded them from the following analysis for the sake of simplicity. Readout weights were averaged over these encoding assemblies and their time evolution during learning is shown in Fig. 3b. Through learning, the neural ensemble encoding a particular chunk developed stronger projections to the corresponding readout unit compared with the other neural ensembles. Consistently with this, the distribution of readout weights was more positively skewed in the encoding ensemble than in the non-encoding ensembles (Fig. 3c). Moreover, a readout unit projected back to the corresponding encoding neuron ensemble more strongly than to the other ensembles (Fig. 3d). Because feedback connections were not modifiable, these results imply that readout connections were strengthened between a readout unit and the reservoir neurons that happened to receive relatively strong feedback from the readout unit. Furthermore, each neural ensemble received slightly stronger inputs from the specific chunk it encoded, which determines the selectivity of the encoding ensemble (Fig. 3e).

**Figure 3.**
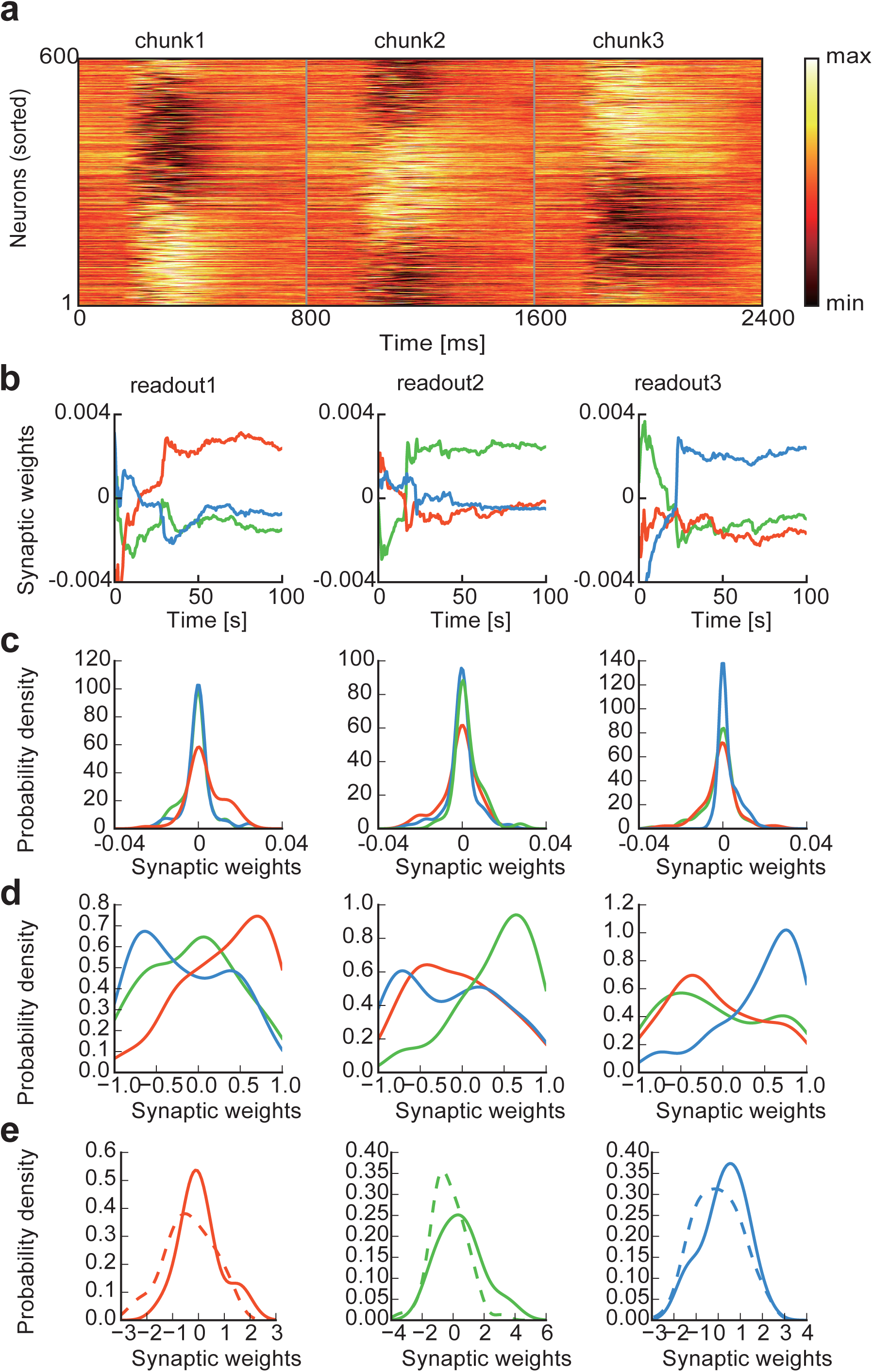
Cell assemblies selected in the reservoirs. **(a)** The activity of each reservoir neuron was averaged over repeated trials and normalized by its maximum activity. Neurons were sorted according to the onset times of their activations to reveal the cell assemblies encoding the three chunks (Methods). **(b)** Temporal evolution is shown for the average weights to the three readout units. **(c)** Normalized distributions are shown for readout weights from each cell assembly. **(d)** The distribution of feedback weights from readout units to each cell assembly is shown. **(e)** The distributions of input weights onto each cell assembly are shown for input neurons belonging to the corresponding chunk (solid) and the others (dashed).

### The role of low-dimensional network dynamics in chunk learning

To get further insight into the mechanism of chunking, we explored the low-dimensional characteristics of the dynamics of reservoir networks. In our model, the two RC modules, say R1 and R2, are thought to mimic others, and this would be possible when the two recurrent networks receiving the same input sequence well predict the responses of other modules. To see how this prediction is formed, we calculated the principal components (PCs) of recurrent network dynamics in the example shown in Fig. 1. After learning, the lowest principal component (PC1) well approximated to the phasic response of the corresponding readout unit during the presentation of chunks (Fig. 4a). Accordingly, the direction of readout weight vector was more strongly correlated with that of PC1 compared to other PCs (Fig. 4b). These results suggest that the low-dimensional characteristics of neural dynamics play a pivotal role in the present chunking.

**Figure 4.**
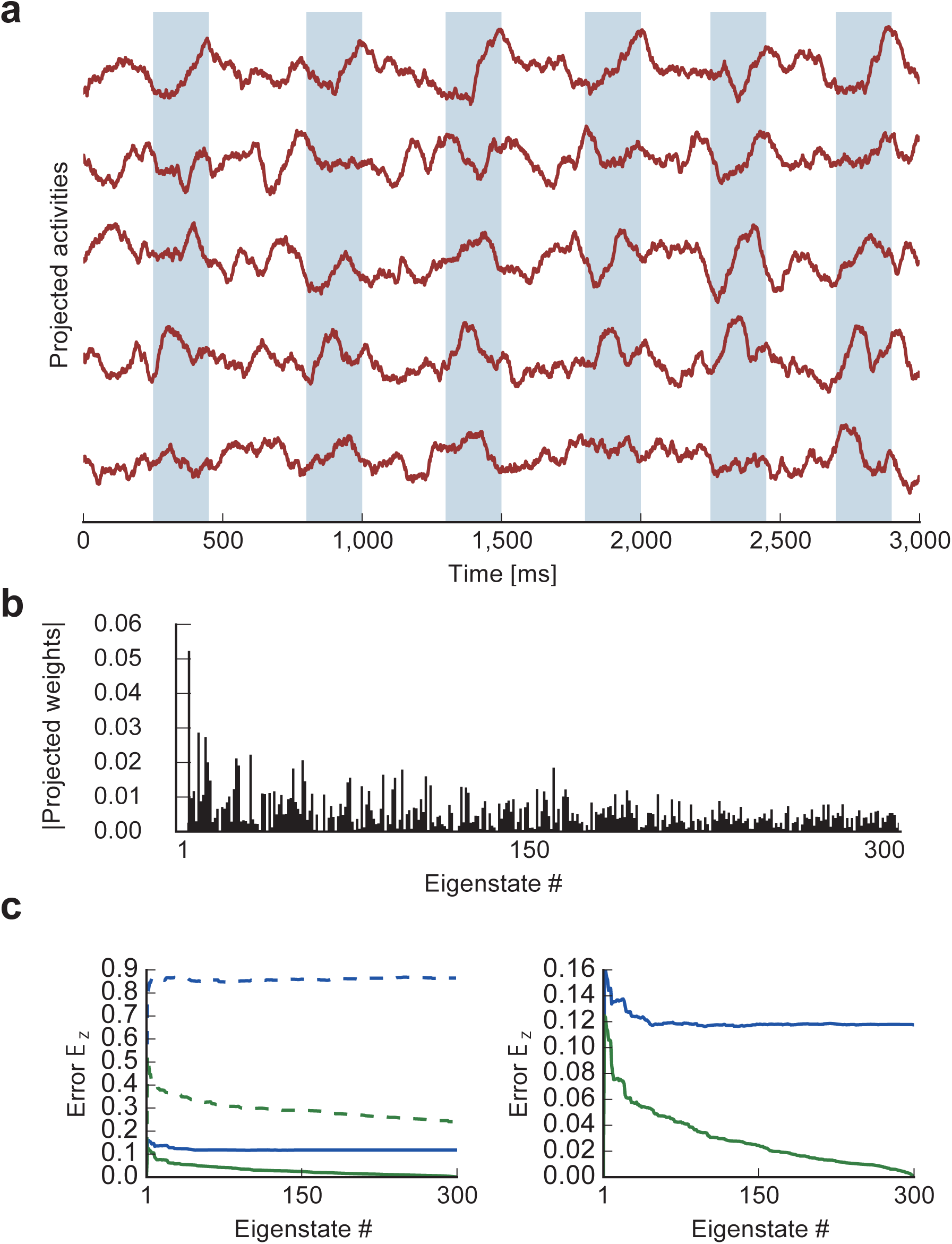
principal component analysis of recurrent networks. Each recurrent network consists of 300 neurons. **(a)** Activities of the reservoir network projected onto the top five eigenvectors of the correlation matrix (PC1 to PC5 from top to bottom) are shown. Shaded areas indicate the intervals of the presentation of chunks. **(b)** Readout weights are projected onto the eigenvectors. **(c)** “Within-self” difference between the R1 and the projected Rl-output (green) and “between-partner” difference between the R2-output and the projected Rl-output (blue) are shown before (dashed) and after (solid) learning.

We then asked to what extent the responses of R1 and R2 are represented by the low-dimensional dynamical characteristics of R1. We calculated the principal components (PCs) of recurrent network dynamics in R1, and expanded its population rate vector and readout weight vector up to the *M*-th order of these PCs (*M*≦*N*_*G*_). Then, we reconstructed the output of R1 by using the *M*-th order rate vector and M-th order weight vector on the low-dimensional subspace spanned by the first *M* PCs (Methods). In Fig. 4c, we calculated differences between the reconstructed R1-output and the full outputs of R1 (within-self difference) and R2 (between-partner difference). Before learning, both differences remained large as *M* was increased. After learning, the “within-self” difference rapidly decreased for *M* < 30-40 and then gradually approached to zero. The “between-partner” difference also rapidly dropped for relatively small values of *M*, but it stooped decreasing for *M* > 50 and remained at relatively large values. These results suggest that R1’s reservoir, and similarly R2’s reservoir, learns to mimic the partner’s response by using the low-dimensional characteristics of its recurrent neural dynamics.

### Network-and chunk-size dependences of learning

Chunk learning may be easier and more accurate if chunks are shorter, but this is not necessarily the case in our model. To see this, we measured learning performance with instantaneous correlations between the activity of a readout unit and a reference response pattern, which takes the value 1 during the presentation of a chunk and 0 otherwise. The correlations were calculated every 15 seconds during learning and averaged over 20 independent simulations. Note that the maximum value of the correlation is 0.5 if the readout activity grows linearly from 0 to 1 during the chunk presentation. Figure 5a shows the correlations for input sequences containing a short or a long chunk (the length 4 or 7, respectively) in the networks of various sizes (*N* = 30, 300 and 500). The correlations were nearly zero before learning and reached to similar maximum values approximately within ten learning steps. The average values of the correlations were generally larger for chunk size 4 than for chunk size 7, but the differences were not statistically significant for any network size tested here (Fig. 5b). These results suggest that learning performance does not significantly depend on the chunk size.

**Figure 5.**
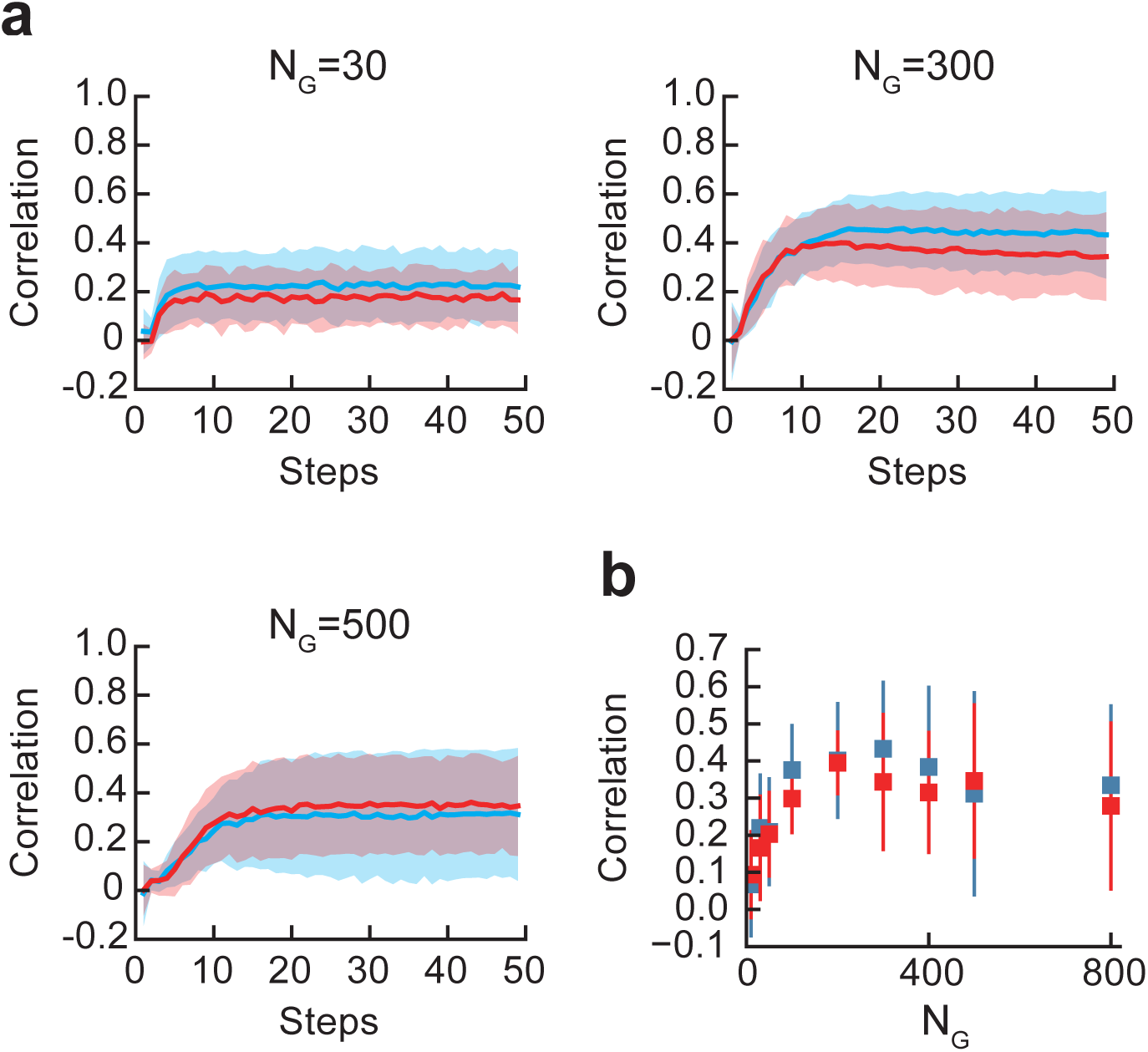
Learning with different sizes of reservoirs and chunks. **(a)** The time course and performance of learning are shown for an input sequence involving a single chunk of the length 4 (blue) or 7 (red). Three networks with different sizes (*N*_G_ = 30, 300, 500) were tested. **(b)** The correlation after learning are plotted as a function of the network size.

In addition, a larger network did not necessarily show better performance. The magnitude of post-learning instantaneous correlations was not significantly increased when the network size was 200 or greater (Fig. 5b). Thus, the performance of chunk learning does not scale with the network size. This seems to be reasonable because increasing the size of reservoirs does not necessarily increase the variety of neural responses useful for learning if the size is already sufficiently large.

### Crucial role of noise in chunk learning

We found that external noise plays an active role in successful chunking. We demonstrate this in the case where input only contains a single chunk (Fig. 6a). In the absence of noise readout units still exhibited phasic responses, but these responses were not necessarily time-locked to chunks. As shown later, the two RC modules in principle may agree on an arbitrary feature of input sequence, which implies the RC system may converge to a local minimum of learning. Noise may help the system to escape from the local minima. On the other hand, too strong noise completely deteriorated the phasic responses to chunks. Thus, the RC system could learn chunks only when a modest amount of external noise existed (Fig. 6b). In the presence of adequate noise (*σ* = 0.25), the average weight of readout connections was rapidly decreased to a small equilibrium value during learning (Fig. 6c), leaving some readout weights much stronger than the majority (Fig. 6d). This reduction was expected because external noise gives a regularization effect on synaptic weights in error-minimization learning (Bishop, 1995). The strong weights were attained for readout connections from the reservoir neurons responding to the chunk, hence were crucial for the chunk detection. However, this was not the case in the absence of noise (*σ* = 0). We counted the fraction of strong readout connections emergent from the chunk-encoding reservoir neurons, where strong connections were such connections that were greater than the standard deviation of the weight distribution. Such a fraction was significantly larger in the presence of adequate noise than in the absence of noise. Under strong noise (*σ* = 1), although the weight distribution becomes more bimodal, noise disrupted learning and the system failed to capture the chunks (Fig. 6e).

**Figure 6.**
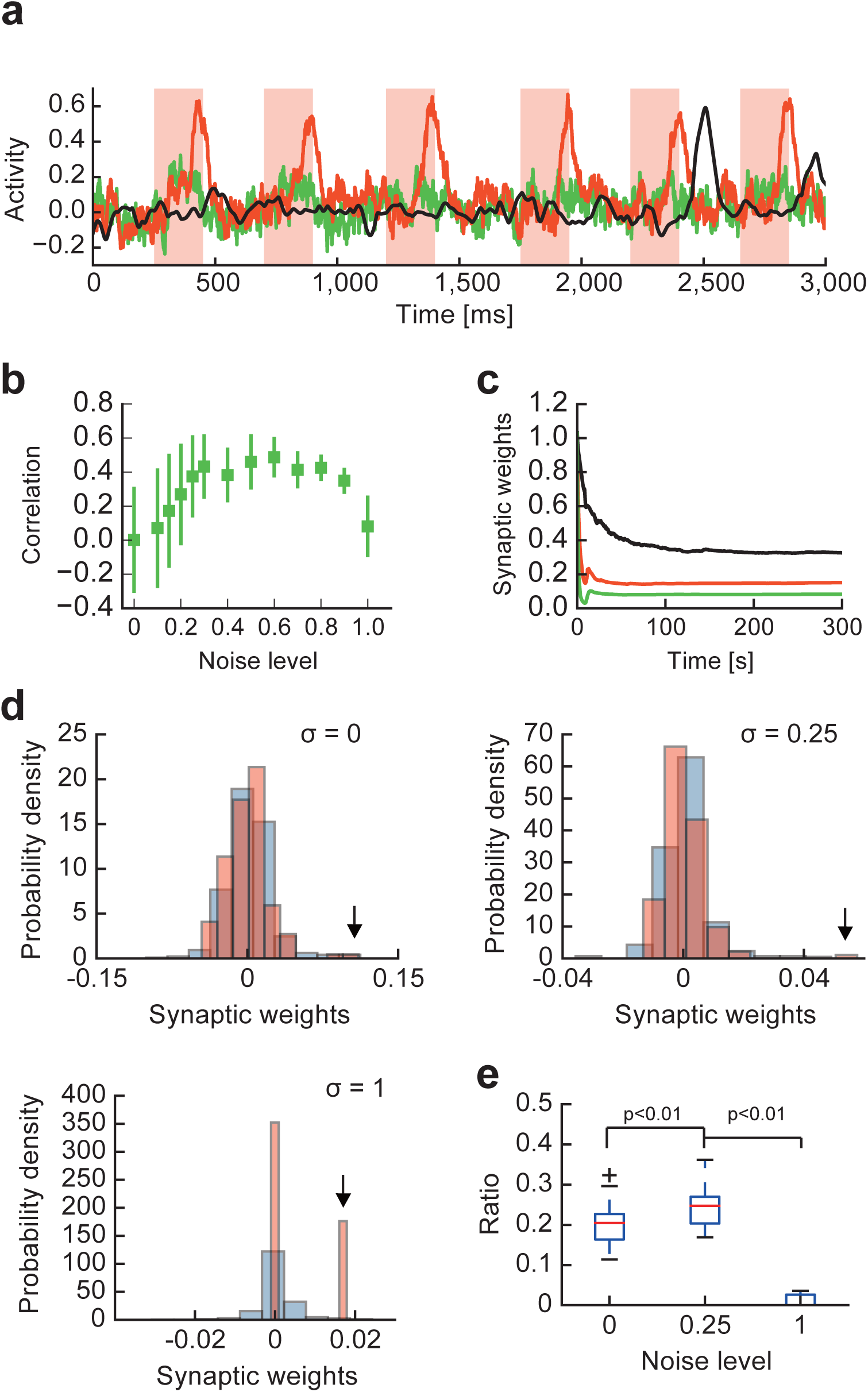
Effects of noise on successful chunk learning. **(a)** Activity of a readout unit after learning a chunk at different noise levels: *σ* = 0 (black), 0.25 (red) and 1 (green). **(b)** Learning performance is a non-monotonic function of the noise level. The optimal performance was shown at *σ* = 0.5~0.6. **(c)** Evolution of the norm of readout weights during learning is shown for *σ* = 0 (black), 0.25 (red) and 1 (green). **(d)** The distributions of readout weights from chunk-encoding (red) and non-encoding (blue) reservoir neurons are shown after learning at different noise levels. Arrows indicate the maximum weight values from the chunk-encoding neurons. **(e)** The fraction of strong readout weights (see the main text) from the encoding neurons is shown for different noise levels. The fraction is significantly larger for *σ* = 0.25 compared with *σ* = 0 and 1 (p<0.01, Mann-Whitney U test).

Though the results so far suggest that mutual supervision enables the RC system to learn the recurring groups of items in a sequence, these results do not indicate how the system chooses the particular groups for learning. Then, the question arises whether our model detects any “chunk” if a sequence merely repeats each alphabet randomly without temporal grouping. To study this, we constructed a set of input sequences of ten alphabets, where all these alphabets appeared equally often in each sequence, and exposed the RC system with a readout unit to these sequences. We found that the system learned to respond to one of the alphabets with approximately equal probabilities (Supplementary Fig. 2a). Then, we made the occurrence probability of alphabet “a” twice as large as the occurrence probabilities of the others and found that the system detected “a” about twice as frequent as the others (Supplementary Fig. 2b). These results indicate that the RC system learns a repeated feature with the probability proportional to its occurrence frequency.

This frequency dependence of our model partially accounts for the features of sequence that are grouped into chunks. Further, as demonstrated in Fig.4 our model engages a pair of RC modules in the mutual prediction of partners’ responses, and this prediction would be easier for the items in input that repeatedly occur in a fixed temporal order. However, the explicit role of temporal grouping in chunking remains to be further clarified.

Finally, we demonstrate that the RC system can simultaneously chunk multiple sequences with overlaps, where input sequences share some alphabets as common items. In some sequences, common alphabets appeared in the beginning or the end of chunks (Fig. 7a), whereas other sequences involve common alphabets in the middle of chunks (Fig. 7d). In both cases, the RC system (with two readout units) successfully chunked these input sequences without much difficulty (Figs. 7b, e). Interestingly, the activity of readout units averaged over repetitive presentations ceased to increase during the presentation of the overlapping part of chunks (Fig. 7c, f). This seems reasonable as overlapping part does not contribute to the prediction of the following items in the chunks and hence needs not be learned.

**Figure 7.**
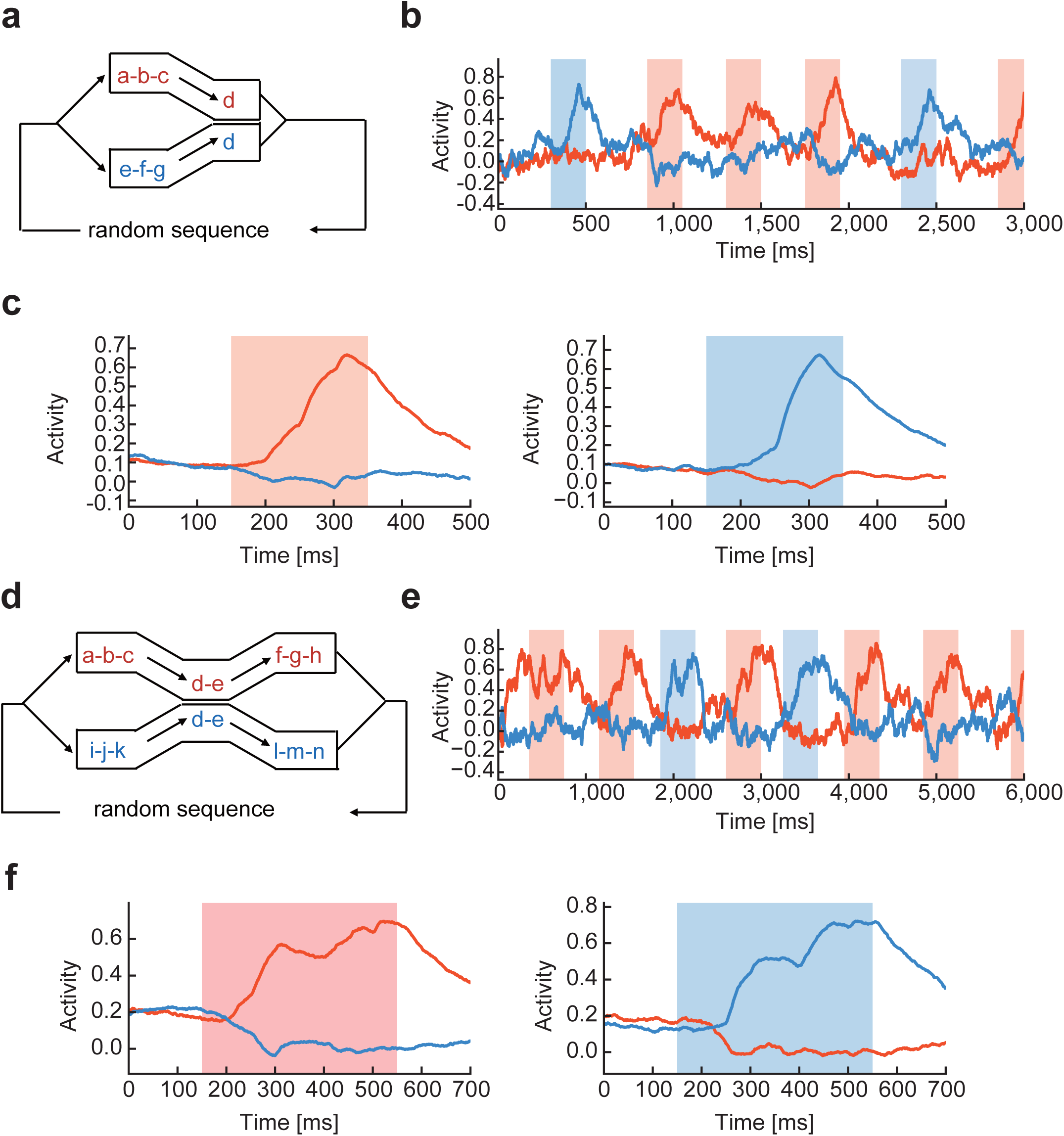
Learning chunks with mutual overlaps. **(a)** Two chunks shared the last component “d” in a random input sequence. **(b)** Activities of two readout units were selective to different chunks after learning. **(c)** The average response profiles are shown for the two readout units. **(d)** Two chunks shared the middle components “d-e” in a random input sequence. **(e) and (f)**, Activities of two readout units and the average response profiles are shown, respectively.

## Discussion

Chunking refers to the segmentation of a continuous flow of stimuli into discrete, temporally bound events. We constructed a dual RC framework for unsupervised learning of multiple chunks hidden in random sequences of alphabets. While each RC system obeys supervised learning (i.e., FORCE learning), the entire system performs unsupervised learning through mutual supervision between the two RC systems. Chunks are represented by, and the probability of detection is proportional to the frequency of repetition in the sequence. The dual RC system successfully detects the groups of alphabets that repeatedly appear in a sequence without a priori knowledge on the contents, lengths and temporal positions of target chunks.

Chunking has been often accounted for by predictive uncertainty or surprise (Reynolds et al., 2007; Baldwin et al., 2008; Zacks et al., 2011; Saffran et al., 1996). This explanation essentially relies on the detection of differences in transition probabilities between different sequence components, i.e., the non-uniform statistical structure of sequence. However, some recent evidence suggests the existence of an alternative mechanism of chunking in which events are segmented on the basis of the temporal community structure of sequential stimuli (Schapiro et al., 2013). It has been shown that individual items in a sequence are concatenated into an event when they frequently go together in the sequence. Our dual RC system automatically chunks a continuous flow of stimuli based on the temporal clustering structure and occurrence probabilities of stimuli without relying on predictive uncertainty or surprise. In this sense, our model supports a neural mechanism relying on the community structure of sequences.

The dual RC system shows good performance in the presence of external noise. Without noise, the system also learns certain segments of sequence, but these segments do not often coincide with any of the repeated chunks. An adequate amount of external noise eliminates such spurious responses and enables the system to respond to the repeated chunks. This finding is interesting because the initial state of the dual RC system is chosen on the so-called “edge of chaos”, on which weakly chaotic neural dynamics provides an adequate amount of flexibility for supervised learning in the RC system (Sussillo and Abbott, 2009; Toyoizumi and Abbott, 2011; Rivkind and Barak 2017). The present system assumes a similar initial state and further requires the regularization of synaptic weights dynamics by noise in addition to the chaotic state evolution (Fig. 6c). A RC system (with a single reservoir) was previously used to account for the observed variety of task-related activity in the motor cortex, and the regularization of synaptic weights greatly contributed to the elimination of strange neuronal responses that were never observed in experiment and were presumably unrelated to the actual motor control (Sussillo et al., 2015). External noise plays a similar role in the present mutually supervising RC system.

It is worthwhile pointing out an interesting similarity between the responses of readout units in our model and those observed in the basal ganglia during the formation of motor habits. If animals repeatedly perform a sequential motor behavior, the behavior becomes more rigid and automatic over the course of learning and practice. This process is accompanied by the chunking of motor sequences, and the basal ganglia is thought to play a pivotal role in habit formation in various species and behaviors (Smith and Graybiel, 2013; Graybiel and Grafton, 2015). For instance, in the rats running on a T maze, the majority of dorsolateral striatal neurons exhibit burst firing when the run is initiated or completed, or both (Barnes et al., 2011). In the mice trained to generate a sequence of actions, an increased population of striatal neurons selectively responds to the initial (Start cells) or the last (Stop cells), or both, action in the sequence (Jin and Costa, 2010; Jin et al., 2014). In our model, readout units always respond strongly to the last component of each chunk, like the Stop cells. On the other hand, our model does not show Start cell-like responses, which is reasonable as chunks are preceded by random sequences and nothing predicts the beginnings of chunks. It is intriguing to investigate whether and how Start cells are formed in the proposed framework of unsupervised learning.

The proposed learning scheme works most efficiently when two RC systems are not interconnected and work independently. A small amount of recurrent connections between them abolishes the ability of chunking. For instance, the whole network could not learn a single chunk if only 60 connections are induced between the two reservoirs each of which has more than 90000 internal recurrent connections. Where can such independent networks located in the brain? Because they are functionally equivalent, it is unlikely that they are implemented in functionally distinct areas. One possibility is that they are represented by mutually disconnected recurrent neuronal networks in a local cortical area. This may occur if the two networks are spatially separated with a sufficiently long distance. Another intriguing possibility is that they are distributed to functionally similar cortical areas in different hemispheres. For instance, inferior frontal gyrus and anterior insula are bilaterally activated when a sequence of visual stimuli is chunked (Bor et al., 2003, Schapiro et al., 2013). Interhemispheric and subcortical projections of cortical pyramidal cells have been extensively studied. Whether subnetworks of pyramidal cells perform chunking in the above or other cortical areas (Zacks et al., 2001) is an intriguing open question.

In sum, we proposed an unsupervised learning system by combining two independent reservoir computing modules. During learning the two systems supervise each other to generate coincident outputs, which in turn allows the entire system to consistently learn chunks hidden in irregular input sequences. As chunking is a fundamental step in the analysis of sequence information, our results have significant implications for understanding how the brain models the external world.

## METHODS

### Neural network model

In this study, the proposed model is composed of two recurrent networks (reservoirs). Each recurrent network is composed of *N*_*G*_ neurons. Each neuron follows the following dynamics as *i* = 1,2,… *N*_*G*_,

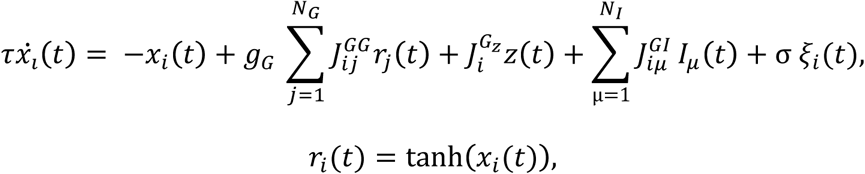

where *ξ*_*i*_(*t*) is a random (Wiener) process and *σ* is the standard deviation. *N*_*I*_ is the number of input neurons. The parameter *g*_*G*_ determines the complexity of the behavior of the reservoir, and shows chaotic spontaneous activity if *g*_*G*_ > 1. The instantaneous output is given by where ***w*** is the readout weight vector. The readout unit is connected with *n* reservoir neurons by the readout weights ***w***. The initial values of the readout weights ***w*** are generated by a Gaussian distribution with the mean 0 and variance 1/*n*. The readout weights are modified according to the FORCE learning rule in which the error between the actual output and the teaching signal is minimized (Sussilo and Abbott, 2009). The activity of the readout unit is transmitted to the reservoir via the feedback. Each element of the feedback coupling *J*^*G*_*Z*_^ is randomly sampled from a uniform distribution [-1, +1]. *J*^*GG*^ is the connection matrix of the reservoir and each element is taken from a Gaussian distribution with mean 0 and variance 1/(*pN*_*G*_), where *p* is the connection probability of the reservoir neurons. In addition, *J*^*GI*^ is the connection matrix between input neurons and the reservoir, and each row has only one non-zero element drawn from a normal distribution of mean 0 and variance 1. We simulated the model with time steps of 1 [ms].

The values of parameters used in simulations are as follows: in Figs. 1, 4 and 6, *N*_*G*_ = 300, *p* = 1,*n* = 300 and *σ* = 0.3; in Figs. 2 and 3, *N*_*G*_ = 600, *p* = 0.5, *n* = 300 and *σ* = 0.3; in Fig. 5, *p* = 1, *σ* = 0.3, *n* = *N*_*G*_, while the values of *N*_*G*_ were varied; in Fig. 7, *p* = 1, *n =* 300, and *N*_*G*_ = 800, *σ* = 0.15 (b) or *N*_*G*_ = 500, *σ* = 0.1 (e). In all simulations, *τ* = 10 [ms], *N*_*I*_ = 26, and *g*_*G*_ = 1.5. The learning rate was set as *α* = 100 because larger values could cause instability in the learning process. The network was trained typically for several hundreds of seconds except in Figs. 2, 7b and 7e where the simulation time was 5000, 2500 and 25,000 [s], respectively.

### Normalized output for teaching signals

In our learning rule, we changed the outputs of readout units such that the mean outputs coincide with zero and the standard deviation becomes unity:

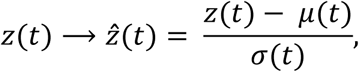

where *μ*(*t*) and *σ*(*t*) were calculated as

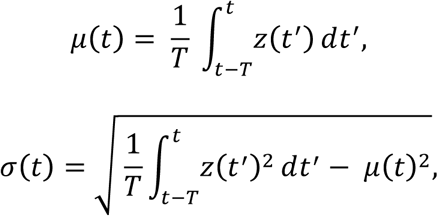

with a sufficiently long period *T* (= 15 [s]). The modified output 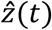 was then transformed by two nonlinear functions to generate the teaching signal shown in the Results.

### Selectivity of reservoir neurons

In Fig. 3a, the activities of all reservoir neurons were first averaged and then normalized. To define the response selectivity of neurons, we sorted all of the neurons by their mean activation phases defined as,

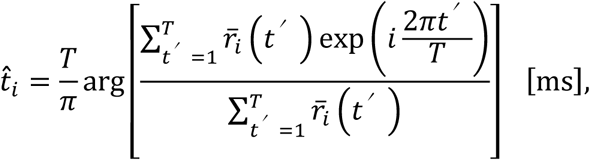

where 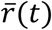 is the normalized average response of each cell and *T* = 2400 [ms]. Each reservoir neuron generally showed a significantly large and prolonged phasic response to a particular chunk, which determined the selectivity of the reservoir neuron. We defined a phasic response as such transient activity that exceeded the threshold value *μ* + 3*σ* for more than 100 [ms], where μ and σ stand for the average and standard deviation of its activity during the presentation of input sequence. Neurons that were not related to any chunks or responded to multiple chunks were discarded in the analysis.

### Analysis of the low-dimensional dynamics of reservoirs

In Fig.4, we expanded the neural responses ***r***_R1_(*t*) of recurrent network in R1 in terms of its top *M*(≦*N*_*G*_) principal components:

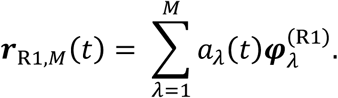

Here, 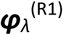 is the *λ*-th eigenvector of R1 reservoir and the coefficient is given as the inner product 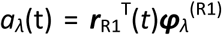. Similarly, we expanded the readout weight vectors from R1 and R2 as

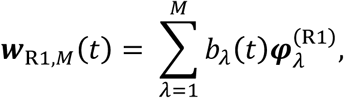

where 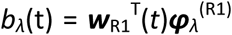. The above expansions represent neural responses and weight vectors projected onto the low-dimensional subspace spanned by the first *M* PCs. We then calculated difference between the actual output of R1 and the output reconstructed on the subspace as

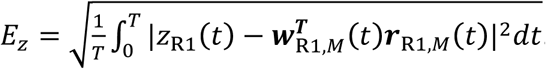

Difference between the output of R2 and the projected R1-output, 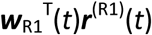, was calculated in a similar fashion.

## Acknowledgements

This work was partly supported by KAKENHI (nos. 15H04265, 16H01289 and 17H06036) to T.F. We thank Vladimir Klinshov, Tomoki Kurikawa, Keita Watanabe and Takashi Takekawa for illuminating discussions about analysis methods for population neural dynamics.

